# Modtector: Ultra-Fast Modification Signal Mining on Mapped Sequencing Reads

**DOI:** 10.1101/2025.10.08.681300

**Authors:** Tong Zhou, Yifan Hong, Panfeng Li, Xitong Liu, Ang Li, Lei Sun

## Abstract

**Summary:** Current tools for RNA epitranscriptomic modification and structural feature analysis are often fragmented, focusing on single signal types and struggling with processing efficiency, especially as sequencing data volumes increase and single-cell technologies advance. To address this challenge, we developed Modtector, an efficient and versatile tool for unified extraction of modification signals from aligned sequencing data. By integrating dual-signal recognition within a single framework, Modtector employs a “count-then-correct” approach to handle both mutation and stop signals, significantly reducing computational complexity. This results in a multi-fold performance improvement compared to existing tools, particularly when processing large-genome, high-coverage datasets—such as completing the analysis of HEK293 22G data in 5 minutes— demonstrating its potential for large-scale data analysis.

**Availability and implementation:** The manual is available at Readthedocs (https://modtector.readthedocs.io/), and source code is available at GitHub (https://github.com/TongZhou2017/modtector) and Crate.io (https://crates.io/crates/modtector).

**Graphic Abstract:** 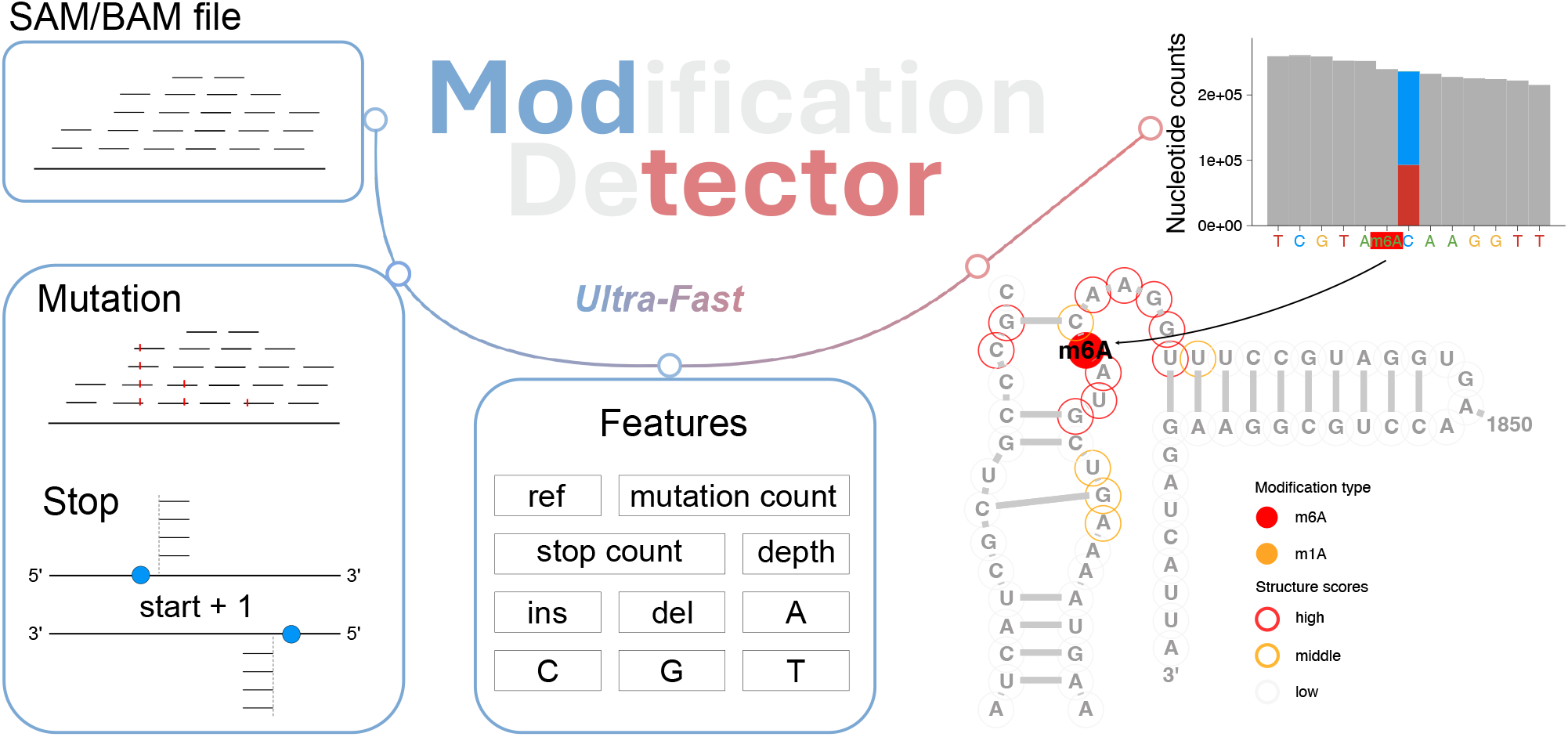

## 1. Introduction

Chemical probing techniques are crucial for studying biomolecular structure and function through high-throughput sequencing (Xu *et al*., 2022; Liu *et al*., 2023). While various computational tools exist for analyzing the resulting signals, most are limited in their ability to jointly process multiple signal types, focusing exclusively on either mutation patterns (Busan and Weeks, 2018) or termination events (Li *et al*., 2020), requiring researchers to use separate tools and manually integrate the results—leading to potential inconsistencies and computational overhead.

This fragmentation becomes particularly problematic when analyzing complex modification patterns involving both mutation and stop signals, such as those seen with RNA structure probing (Sexton *et al*., 2017) and m6A modifications (Liu *et al*., 2023). The absence of unified frameworks complicates analytical workflows and hinders direct comparisons between different signal types under consistent statistical conditions (Sexton *et al*., 2017). Furthermore, as sequencing data grows exponentially, computational efficiency has become a key consideration for large-scale analyses (Mu *et al*., 2025). For instance, RASP, which collects over a thousand datasets, struggles to perform genome-wide reanalysis within a reasonable time due to the inefficiency of existing tools (Li *et al*., 2021). Similarly, single-cell data—often generating large volumes of independent datasets for multiple cells—presents challenges for tools like bam-readcount, which is slow and can only process mutation signals, limiting the ability to support future stop signal-based single-cell RNA structural analysis (Wang *et al*., 2024).

To address these challenges, we developed **Modtector (Mod**ification de**tector)**, a unified computational framework that simultaneously extracts and correlates multiple modification signals from aligned sequencing reads. By integrating dual-signal quantification within a single statistical model, Modtector enables researchers to obtain a comprehensive view of modification events in a single execution, ensuring consistent analytical conditions across signal types while delivering superior computational performance.

## 2. Features

### 2.1 Unified Framework for Dual-Signal Analysis

Modtector introduces an integrated pipeline (**Figure 1A**) that simultaneously processes two key signal types from SAM/BAM files: nucleotide mismatches (mutation) and reverse transcription truncation events (stop). This unified framework includes core functions such as signal counting, reactivity analysis, normalization, evaluation, and visualization, supporting both primary signal calling and downstream analysis methods. Unlike existing tools that typically require separate analyses for each signal type (**Table S1**), Modtector ensures consistent statistical treatment of both signals by using shared genomic coordinates.

**Figure 1.**
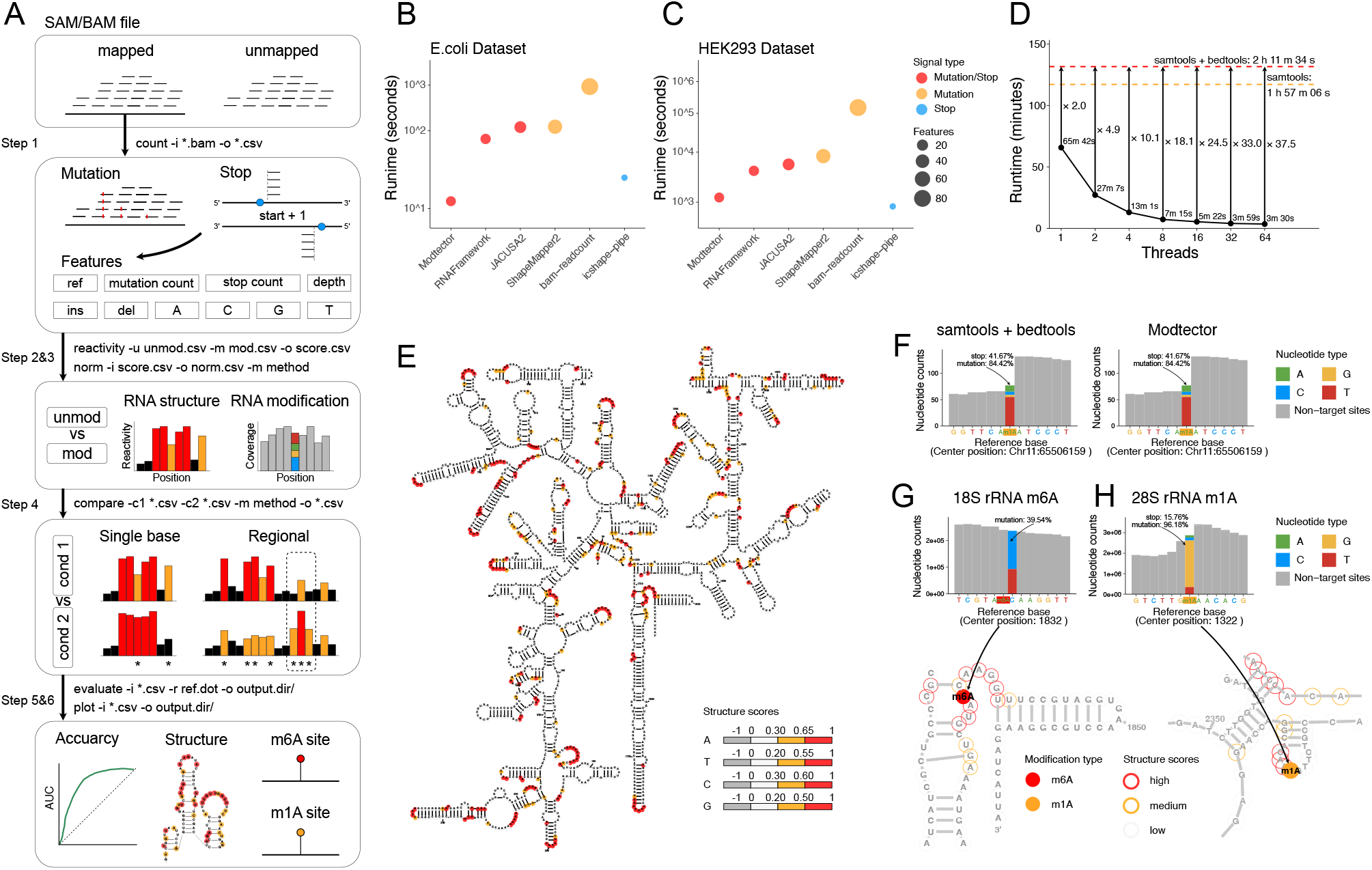
Overview of Modtector’s dual-signal analysis framework and benchmarking results. (A) Workflow of Modtector’s analysis pipeline, illustrating the key steps in processing mapped SAM/BAM files to generate CSV files for mutation and stop signal, applicable to RNA structure and modification data analysis. The flowchart includes command-line examples and key parameters for core functions. (B-C) Runtime comparisons (thread = 1) of Modtector with other tools for RNA structure data analysis using *E. coli* (B) and human HEK293 cell line (C) datasets. Each point represents a tool, with colors indicating the types of signals processed: orange for mutation signals, blue for stop signals, and red for tools supporting both signals. The point size represents the number of features collected. (D) Performance scaling analysis comparing Modtector to samtools and bedtools with varying thread counts (1 to 64). The x-axis represents the number of threads, and the y-axis represents runtime, with log10 scaling applied. Modtector shows significant speed improvements across different thread counts (arrows from data points to the dashed line represent the speedup factor). (E) Visualization of secondary structure of 18S rRNA (icSHAPE dataset: SRR11164863) by Modtector plot command. The reactivity scores calculated by Modtector are overlaid on the structures, with color gradients representing different levels of reactivity: red for high reactivity, yellow for medium reactivity, and grey for low reactivity. (F) Dual-signal distribution of MALAT1 Chr11:65506159 m1A site (m1A-seq dataset: SRR5418425), comparing the consistency between samtools + bedtools and Modtector methods. (G-H) Dual-signal distribution and secondary structure mapping for 18S rRNA 1832 m6A site (G; miCLIP dataset: SRR2003406) and 28S rRNA 1322 m1A site (H; dataset: SRR5418425). The miCLIP method is characterized by mutations or truncations occurring at the previous C position during reverse transcription at m6A sites. The secondary structure backbone downloads from the RNAcentral database (URS0000726FAB_9606 for 18S rRNA, URS000075EC78_9606 for 28S rRNA), and the loop structure scores are mapped through Modtector plot command, with red for high reactivity, yellow for medium reactivity, and grey for low reactivity. Filled circles represent RNA modification sites, with red for m6A modification and orange for m1A modification.

Modtector applies an efficient “count-then-correct” approach to handle stop signal shifts, minimizing redundant file operations. For the core function signal detection (**Figure 1A**, Step 1), Modtector simultaneously quantifies mutation and stop signals across both strands. For stop signal, it first distinguishes strand-specific reads and uses the read start position as a raw signal. Signal shift corrections, such as those needed for RNA structure modifications (e.g., occurring at +1), are applied at the site level. The framework supports both gene-level and genome-level calculations, with genome-wide analysis benefiting from sliding window segmentation and dynamic task deployment to optimize computational resource usage. Overlapping windows (based on stop signal correction lengths) ensure no data loss, addressing issues observed in tools like JACUSA2 (**Figure S1A**).

For other downstream analysis functions, Modtector offers multiple reactivity calculation and data normalization options. It also provides two distinct approaches for both stop (Ding *et al*., 2014; Rouskin *et al*., 2014) and mutation (Siegfried *et al*., 2014; Zubradt *et al*., 2017) signal (**Figure 1A**, Step 2&3). Additionally, Modtector supports differential site statistics at both single-base and region (deltaSHAPE (Smola *et al*., 2015) and Diffscan (Yu *et al*., 2022)) levels for different sample groups (**Figure 1A**, Step 4). The evaluate function assesses accuracy among base type seperately, providing multiple performance metrics (eg. AUC, F1-score) to ensure the reliability of the extracted signals (**Figure 1A**, Step 5). For the signal score and RNA modification site annotation, Modtector can visualize in secondary structure in SVG format (**Figure 1A**, Step 6).

### 2.2 Enhanced Performance Compared to State-of-the-Art Tools

Comprehensive benchmarking comparing Modtector with leading tools (**Table S1**) in RNA structure task, such as RNA Framework (Incarnato *et al*., 2018), JACUSA2 (Piechotta *et al*., 2022), ShapeMapper2 (Busan and Weeks, 2018), bam-readcount (Khanna *et al*., 2022), and icshape-pipe (Li *et al*., 2020), reveals significant runtime advantages on *E. coli small genome* and human datasets (**Figure 1B,C**). Notably, it demonstrated a speedup of over 100-fold compared to bam-readcount, which is often used for single-cell data analysis (Wang *et al*., 2024), highlighting Modtector’s potential for large-scale single-cell data processing.

When compared with classic tools in RNA modification task, Modtector achieved a twofold speedup over conventional samtools (Danecek *et al*., 2021) and bedtools (Quinlan and Hall, 2010) pipelines (**Figure 1D**). In typical workflows, samtools/bedtools provide predominantly single-threaded core operations and produce intermediate outputs that require additional, user-written scripts to compute extended features, which limits scalability on large-genome, high-coverage datasets. This constraint also indirectly affects tools that depend on these utilities (e.g., RNA Framework; ShapeMapper2). By contrast, Modtector employs a fully Rust-based, unified dual-signal recognition algorithm and leverages multi-threading throughout, including direct bindings to the htslib crate, thereby maximizing computational efficiency and throughput. For example, using only 8 threads, Modtector can process a 22G human dataset in just 5 minutes, whereas other multi-threaded tools require 64 threads to achieve comparable efficiency (**Table S2**).

### 2.3 Methodological Consistency and Biological Accuracy

In addition to its speed, Modtector demonstrates high consistency with results from other established methods. Most notably, it showed perfect correlation (Pearson’s r = 1.0) with samtools-based approaches for RNA modification data analysis (**Figure S2**), confirming that the computational advantages do not come at the cost of accuracy. This consistency is further evidenced by Modtector’s ability to produce biologically meaningful RNA secondary and structural mappings (**Figure 1E, S3**). For example, base-resolution accuracy assessments revealed higher detection rates for adenosine and cytidine (**Figure S3A**), consistent with DMS-based probing methods (Xu *et al*., 2022).

In addition, Modtector’s RNA modification signal detection is highly consistent, as shown in **Figure 1F**, where it simultaneously identifies dual-signal distributions in the human lncRNA MALAT1 m1A site (Chr11: 65506159). Due to its ability to process both RNA structure and modification signals, Modtector can focus on specific RNA modifications within particular structures, such as the m6A modification on 18S rRNA 1832 (**Figure 1E,G**), which is involved in translation regulation (Sepich-Poore *et al*., 2022), or the m1A modification on 28S rRNA 1322 (**Figure 1H, S3B**), which plays a role in ribosomal structure and function regulation (Sharma *et al*., 2018).

This combination of biological accuracy and methodological consistency makes Modtector a robust and reliable solution for modern chemical probing data analysis.

## 3 Usage and document

Modtector is implemented in Rust and freely available under the MIT License at https://github.com/TongZhou2017/modtector. The software provides a unified command-line interface for simultaneous processing of mutation and stop signals, with subcommands for specialized tasks. Designed for Unix-like environments, the binary executable of Modtector can be directly copied and used without the need for installation, making it easy to integrate into existing bioinformatics workflows. Comprehensive documentation, example datasets, and benchmarking scripts are included to facilitate adoption and reproducibility at readthedocs website (https://modtector.readthedocs.io/).

## Acknowledgements

We would like to thank Yongkang Tang and Ruobin Zhao for valuable suggestions on functionality, Shaozhen Yin for assistance with accuracy assessments, and Yuning Liu for testing the code and providing insightful feedback. Special thanks to the Computing Platform at the Core Facility and Service Platform, School of Life Sciences, Shandong University, for providing computational resources and support.

## Data availability

All data are incorporated into the article and its online supplementary material.

## Figure Legends

**Figure S1.**
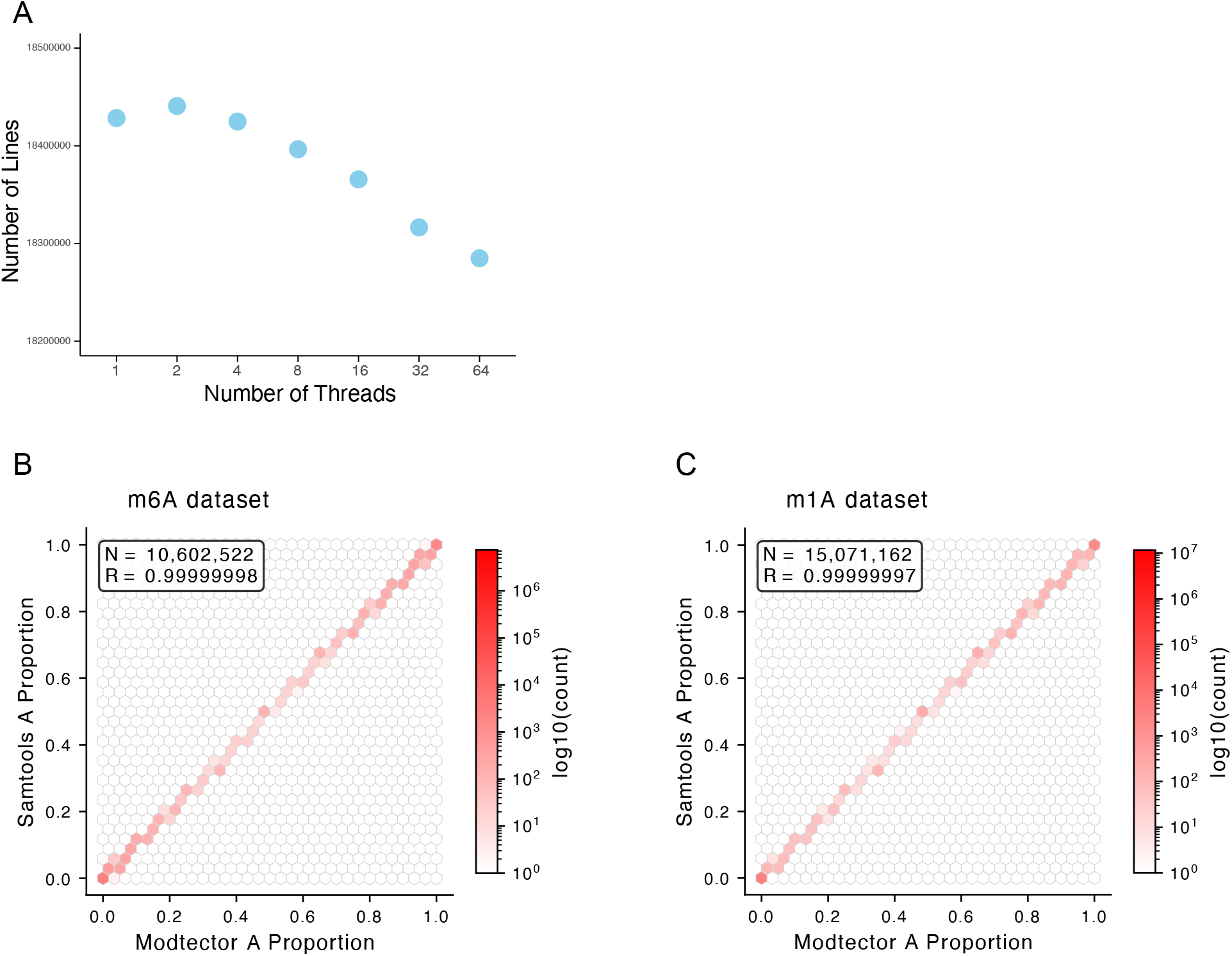
Result consistency comparison. (A) JACUSA2 loses data in multi-thread models due to overlapping signal corrections. The x-axis represents the number of threads (1 to 64), and the y-axis represents the number of output file rows. (B-C) Pearson’s correlation analysis between Modtector and samtools-based m6A (B) and m1A (C) RNA modification data. The x-axis represents the proportion of A bases at genomic single-base sites for Modtector, and the y-axis represents the results from samtools. The dark of the hexagonal color blocks represents site density.

**Figure S2.**
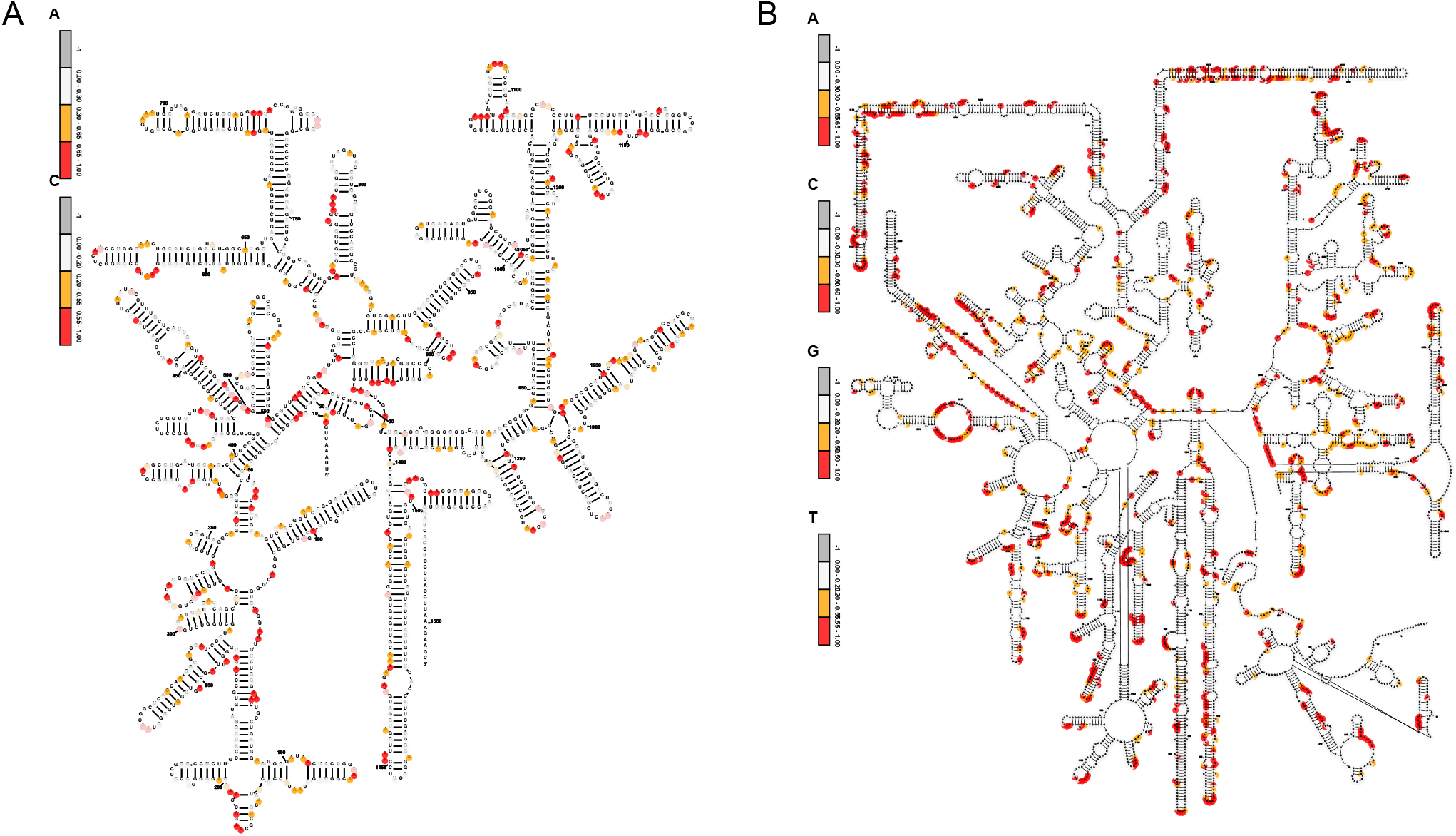
Visualization of secondary structure of RNA by Modtector plot command. (A-B) E.coli 16S rRNA (DMS-seq dataset: SRR3710405) and human 28S rRNA (icSHAPE dataset: SRR11164863) reactivity scores calculated by Modtector are overlaid on the structures, with color gradients representing different levels of reactivity: red for high reactivity, yellow for medium reactivity, and white for low reactivity.

## Table Legends

**Table S1. List of tools for benchmark comparison**.

**Table S2. Runtime performance in multi-thread model**.

## References

Busan, S. and Weeks, K.M. (2018) Accurate detection of chemical modifications in RNA by mutational profiling (MaP) with ShapeMapper 2. RNA, 24, 143–148.

Danecek, P. et al. (2021) Twelve years of SAMtools and BCFtools. GigaScience, 10, giab008.

Ding, Y. et al. (2014) In vivo genome-wide profiling of RNA secondary structure reveals novel regulatory features. Nature, 505, 696–700.

Incarnato, D. et al. (2018) RNA Framework: an all-in-one toolkit for the analysis of RNA structures and post-transcriptional modifications. Nucleic Acids Research, 46, e97–e97.

Khanna, A. et al. (2022) Bam-readcount-rapid generation of basepair-resolution sequence metrics. JOSS, 7, 3722.

Li, P. et al. (2020) icSHAPE-pipe: A comprehensive toolkit for icSHAPE data analysis and evaluation. Methods, 178, 96–103.

Li, P. et al. (2021) RASP: an atlas of transcriptome-wide RNA secondary structure probing data. Nucleic Acids Research, 49, D183–D191.

Liu, C. et al. (2023) Absolute quantification of single-base m6A methylation in the mammalian transcriptome using GLORI. Nat Biotechnol, 41, 355–366.

Mu, K. et al. (2025) RASP v2.0: an updated atlas for RNA structure probing data. Nucleic Acids Research, 53, D211–D219.

Piechotta, M. et al. (2022) RNA modification mapping with JACUSA2. Genome Biol, 23, 115.

Quinlan, A.R. and Hall, I.M. (2010) BEDTools: a flexible suite of utilities for comparing genomic features. Bioinformatics, 26, 841–842.

Rouskin, S. et al. (2014) Genome-wide probing of RNA structure reveals active unfolding of mRNA structures in vivo. Nature, 505, 701–705.

Sepich-Poore, C. et al. (2022) The METTL5-TRMT112 N6-methyladenosine methyltransferase complex regulates mRNA translation via 18S rRNA methylation. Journal of Biological Chemistry, 298, 101590.

Sexton, A.N. et al. (2017) Interpreting Reverse Transcriptase Termination and Mutation Events for Greater Insight into the Chemical Probing of RNA. Biochemistry, 56, 4713–4721.

Sharma, S. et al. (2018) A single N1-methyladenosine on the large ribosomal subunit rRNA impacts locally its structure and the translation of key metabolic enzymes. Sci Rep, 8, 11904.

Siegfried, N.A. et al. (2014) RNA motif discovery by SHAPE and mutational profiling (SHAPE-MaP). Nat Methods, 11, 959–965.

Smola, M.J. et al. (2015) Detection of RNA–Protein Interactions in Living Cells with SHAPE. Biochemistry, 54, 6867–6875.

Wang, J. et al. (2024) RNA structure profiling at single-cell resolution reveals new determinants of cell identity. Nat Methods, 21, 411–422.

Xu, B. et al. (2022) Recent advances in RNA structurome. Sci. China Life Sci., 65, 1285–1324.

Yu, B. et al. (2022) Differential analysis of RNA structure probing experiments at nucleotide resolution: uncovering regulatory functions of RNA structure. Nat Commun, 13, 4227.

Zubradt, M. et al. (2017) DMS-MaPseq for genome-wide or targeted RNA structure probing in vivo. Nat Methods, 14, 75–82.

